# Local gradient analysis of human brain function using the Vogt-Bailey Index

**DOI:** 10.1101/2022.10.14.511925

**Authors:** Christine Farrugia, Paola Galdi, Irati Arenzana Irazu, Kenneth Scerri, Claude J. Bajada

## Abstract

In this work, we take a closer look at the Vogt-Bailey (VB) index, proposed in Ref. [1] as a tool for studying local functional homogeneity in the human cortex. We interpret the VB index in terms of the minimum ratio cut, a parameter that indicates whether a network can easily be disconnected into two parts having a comparable number of nodes. In our case, the nodes of the network consist of a brain vertex/voxel and its nearest neighbours, and a given edge is weighted according to the affinity of the nodes it connects (as reflected by the modified Pearson correlation between their fMRI time series). Consequently, the minimum ratio cut quantifies the degree of small-scale similarity in brain activity: the greater the similarity, the ‘heavier’ the edges and the more difficult it is to disconnect the network, hence the higher the value of the minimum ratio cut. We compare the performance of the VB index with that of the Regional Homogeneity (ReHo) algorithm, commonly used to assess whether voxels in close proximity have synchronised fMRI signals, and find that the VB index is uniquely placed to detect sharp changes in the (local) functional organization of the human cortex.

## 1 Introduction

Since the emergence of neuroscience as a distinct discipline in the 1950s and early 1960s, there has been increasing interest in understanding the organizational principles of the cerebral cortex. The degree to which the cortex is parcellated into separate regions has been strongly debated over the years. Oskar and Cécile Vogt, well-known for their myeloarchitectonic cortical maps, counselled sharp, ‘hairline’ boundaries ([2] and references therein). The cytoarchitectonic parcellation described by Korbinian Brodmann [3] also divides the cortex into different areas, although Brodmann himself pointed out that in some cases the boundaries are not sharp and changes occur gradually [3]. On the other hand, Percival Bailey and Gerhardt von Bonin advocated that the isocortex is characterised by a high degree of homogeneity [4]. This raises the question: is it really appropriate to parcellate the cortex into distinct regions?

The rapid advance of new technologies and introduction of techniques such as magnetic resonance imaging (MRI), coupled with developments in the fields of graph theory and network analysis, gave new impetus to the study of cortical organization. The aim of applying graph theory to neural data is to investigate the emerging connectivity patterns, which reveal how different brain areas are related to each other structurally or functionally [5]. The use of networks in neuroscience can provide important insight into human cognition and behaviour [6], [7], and further our understanding of how the brain changes with age [8]–[10], how it adapts itself to various cognitive demands [11], and how intelligence and intellectual abilities are related to functional connectivity [12], [13]. Functional networks have also been widely studied in the context of neurological and psychiatric disorders such as the degenerative dementias [14], epilepsy [15], multiple sclerosis [16], schizophrenia [17], depression [18] and autism spectrum disorder [19]. In recent years, the concept of *gradients* (or spatial transitions) in brain organization [20] has become an especially popular topic. These gradients are usually used to infer how the ‘activation hubs’ for a certain class of functions are distributed within the brain.

In the majority of cases, neuroscientific works consider connections between different brain regions (or ‘parcels’). However, methods from graph theory can also quantify the affinity between two or more adjacent vertices/voxels. This property makes them a good tool to study the local disruptions in brain function implicated in certain disorders/diseases [21]–[23]. The searchlight Vogt-Bailey (VB) algorithm developed in [1] determines functional connectivity on a per-vertex level by constructing, for each vertex on the surface of the brain, a graph consisting of the original vertex and its neighbours. The vertices are treated as the nodes of the graph, while the (modified) Pearson correlation between their fMRI time series is used to assign weights to the connecting edges. The algebraic connectivity, equivalent to the second smallest eigenvalue of the Laplacian, indicates how easy or difficult it is to disconnect the graph,^1^ and thus serves to gauge the degree of homogeneity around the original vertex. The VB index is defined as a scaled version of the algebraic connectivity. It can additionally be adapted to serve as a metric for full brain or region-of-interest analysis, depending on the size of the neighbourhood that is provided as input. The VB toolbox is available at https://github.com/VBIndex.

The searchlight VB index has been used (albeit in voxel space) to explore the organization of axonal fibres by providing a measure of the correlation between the connection probabilities of neighbouring voxels [24]. The authors modelled the connection probability of a given voxel as a data series consisting of 360 elements, with each element reflecting the probability of tract linkage between the voxel in question and one of 360 target cortical regions [24]. In another study, the VB index was employed to probe the functional organisation of the rodent hippocampus and indicated a sharp change in connectivity [25].

A number of research articles use the same mathematical basis as the VB index to describe how neural gradients are mapped and how inter-areal boundaries can be identified [26]–[29]. The searchlight VB index differs in that it shifts the focus from the global level to the local – it is calculated by constructing networks on a small scale (one per vertex), rather than across a region of interest or the entire brain.

One shortcoming of local-scale analysis is the artificial enhancement of correlations that may result from volume-to-surface mapping, especially in the vicinity of narrow sulci and gyri [30]. This gets more pronounced when surface resolution is increased with respect to voxel data resolution, since more surface vertices sample the same voxels. To mitigate the issue, another VB index approach was developed – the hybrid searchlight algorithm. This estimates the algebraic connectivity in volumetric space and maps the results to the original surface vertices [30]. A modified version was used with diffusion data to study the impact of preterm birth on the homogeneity of tissue micro structure in the neonatal cortex [31].

The work presented here complements the original article [1] and expounds on the mathematical framework of the VB index. In Section 2, we first take a look at some principles from graph theory, then proceed to an interpretation of the VB index as a cut-set weight corresponding to a cut that partitions the graph into two while attempting to minimize any imbalance in cluster size. Section 3 is dedicated to experimental validation, including comparison with the ReHo metric. We conclude in Section 4.

## 2 Methods

### 2.1 Spectral graph theory: optimisation as an eigenvalue problem

#### 2.1.1 Preliminaries

Let *G* = (*V, E*) be an undirected, weighted graph with vertex set *V* of size *n* and edge set *E* = *{*(*υ*_*i*_, *υ*_*j*_) *∈ V × V, υ*_*i*_ υ_*j*_*}*. G has no loops (i.e. no edges starting and terminating at the same vertex), and direct connections between any two vertices are limited to at most one edge. To simplify notation we shall refer to the edge joining vertices *υ*_*i*_ and *υ*_*j*_, (*υ*_*i*_, *υ*_*j*_), as e_*ij*_. The weight associated with e_*ij*_ will be denoted by w_*ij*_ and is a number in the interval [0, 1].

To determine the best way of partitioning V into two disjoint clusters, we introduce the cost function U:

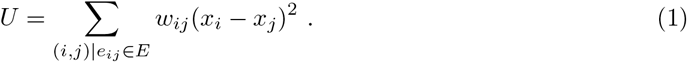

The variables *x*_*i*_ and *x*_*j*_ are the positions of the vertices *υ*_*i*_ and *υ*_*j*_, respectively; we emphasise, however, that they do *not* refer to the positions that the vertices have in 2D Euclidean space (such as in Fig. 2), but rather to coordinates in 1D space. In other words, we map the vertices to a line, much like the beads in one row of an abacus frame, and let the mathematics adjust the vertices (‘beads’) until the separations between them are optimal – in the sense that they make *U* as small as possible. The optimization algorithm looks for a trade-off between the weights of the edges and the separation of the respective vertices. Since we want to minimise *U*, an edge with substantial weight will tend to be shorter, so that the small value of (*x*_*i*_ *− x*_*j*_)^2^ compensates for the large *w*_*ij*_; consequently, in such cases *x*_*i*_ and *x*_*j*_ are usually either both positive or both negative. On the other hand, edges with a relatively small weight can afford to be longer (one might argue that keeping them short would be even better, as it would decrease the cost function further; however, it must be remembered that moving one vertex with respect to another also moves it relative to the remaining vertices, so the question is how to balance edge weights and vertex separations). The end result is that vertices which are strongly connected to each other will usually cluster on one side of the zero reference point when mapped to a line, while vertices sharing a weak connection are further apart and often tend to be positioned on different sides of the zero reference point.

Next, we construct a vector 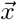 consisting of as many components as there are vertices, with each component *x*_*i*_ being the position of the corresponding vertex *υ*_*i*_ on the line. Since Eq. (1) only constrains the difference between the elements of 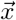, and not the elements per se, it could trivially be satisfied by setting *x*_*i*_ = *x*_*j*_ for all vertices *υ*_*i*_ and *υ*_*j*_. However, condensing all the vertices to one point would hardly constitute a useful solution. We therefore impose the condition [32]

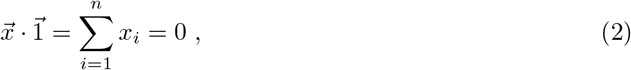

Where 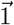 is the all-ones vector of size n. Eq. (2) implies that the components of 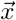 cannot be all positive or all negative, and is essentially an attempt to get well-balanced clusters distributed on each side of the zero reference point.

Let us now take a short detour to define some terminology related to graph theory, starting from the (weighted) *affinity matrix* **A**, whose elements are given by

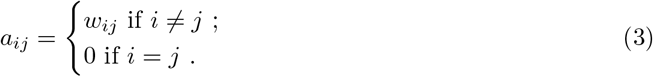

Given that the graph is assumed to be undirected, it follows that *w*_*ij*_ = *w*_*ji*_, and hence **A** is symmetric.

Another array we shall be using is the *degree matrix* **D**. This is a diagonal matrix which may be constructed from **A** by summing its entries either row-wise or column-wise. More specifically,

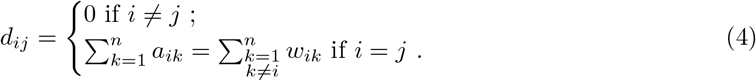

The first equality in the last line follows from Eq. 3, and the quantity 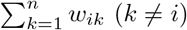 represents a sum over the weights of the edges joining *υ*_*i*_ to the remaining *n* − 1 vertices of the graph (this does not mean the graph is complete; we treat absent edges as edges having a weight of zero). The *i*^th^ element along the principal diagonal of **D**, *d*_*ii*_, is known as the *degree* of the vertex *υ*_*i*_.

The *graph Laplacian* **L** is defined as

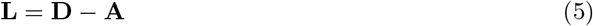

and is a symmetric, positive semi-definite matrix. The spectrum of eigenvalues of the Laplacian can be determined by solving the eigenvalue equation:

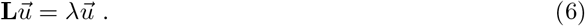

Vectors 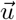 (excluding 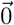, which represents the trivial solution) and values of λ that satisfy this equation are called *eigenvectors* and *eigenvalues*, respectively; each eigenvalue is paired with at least one eigenvector. The graph Laplacian has the smallest eigenvalue equal to 0 and the all-ones vector 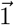 as the corresponding eigenvector. The second smallest eigenvalue is known as the *algebraic connectivity* [33] and is the quantity we will be using to construct the VB Index.

#### 2.1.2 Minimising the Rayleigh quotient

It can be shown that the quadratic cost function of Eq. (1) may be expressed in terms of the Laplacian via the relation ^2^

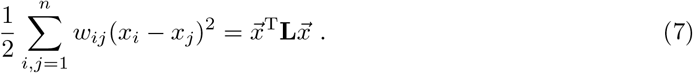

Consequently, the problem of partitioning the graph in the manner outlined above can mathematically be formulated [using Eq. (1)] as the requisite to minimise the Rayleigh quotient:

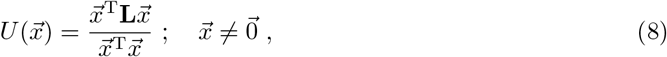

subject to the condition ^3^ 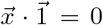 [Eq. (2)]. We have introduced the magnitude of 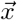 in the denominator to avoid getting solutions which optimise the cost function by making the components of 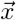 arbitrarily small.

The vector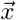 which minimizes 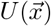 is none other than the Fiedler vector, the eigenvector associated with the second smallest eigenvalue of the Laplacian matrix. This can be proved as follows [35], [36]: let **L** have eigenvalues 0 = λ_1_ < λ_2_ *≤* … *≤* λ_*n*_ with corresponding orthonormal eigenvectors ^4^ 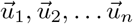. As the eigenvectors provide an orthonormal basis, any vector 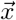 may be expressed in the form

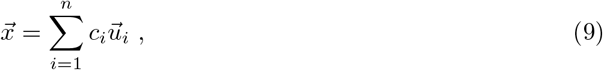

where 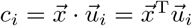. Hence we get that

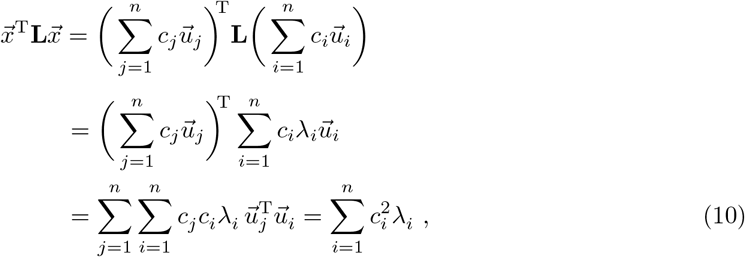

since the eigenvectors are orthonormal (meaning that 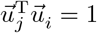 if *i* = *j*, and 0 otherwise). In similar fashion,

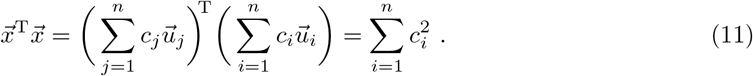

Suppose, now, that 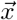 is orthogonal to 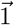, i.e. 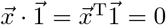. Given that 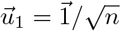, and that the component of 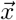 along 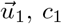, c_1_, is obtained by taking the dot product of 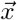 with 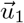, it follows that 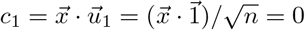. Consequently, we can drop c_1_ from the sums in Eqs. (10) and (11), and write:

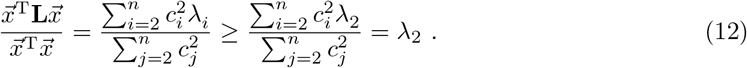

If 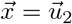,

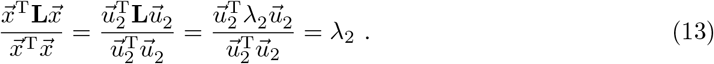

In conclusion, then, any vector 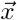 orthogonal to 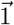 satisfies

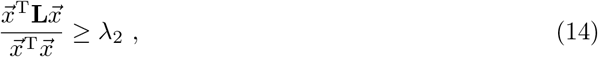

with equality attained when 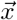 is set to 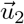, the Fiedler vector. To recap, we have thus far:

- Considered a general vector 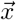 whose elements correspond to the positions of the vertices in 1D space;
- Shown that the partitioning of the graph is optimized [with respect to Eq. (1)] if 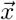 is the Fiedler vector.

In other words, if we would like to separate the graph into two balanced clusters in a way that minimizes the cost function given by Eq. (8), the vertex *υ*_*i*_ should be mapped to the coordinate *x*_*i*_ in 1D space given by the *i*^th^ component of the Fiedler vector. Vertices that end up in close proximity in this 1D setting can be grouped together when the original graph is partitioned.

#### 2.1.3 The generalised vs standard eigenvalue problem

Let us go back to the equation whence the Fiedler vector originated; namely, the standard eigenvalue equation:

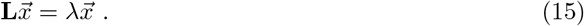

Outlying vertices are likely to have significant effect on the clustering, but we can reduce their influence by using the *generalised eigenvalue problem* in place of the standard one:

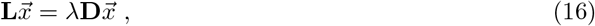

**D** being the degree matrix introduced in Eq. (4). If we now multiply both sides of Eq. (16) by the inverse of the degree matrix, **D**^*−*1^, we get the relation:

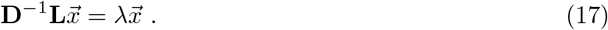

Using the definitions of the degree matrix and the Laplacian from Eqs. (4) and (5), respectively, it is straightforward to show that the matrix product **D**^*−*1^**L** (which is known as the *random walk normalised Laplacian* and will be denoted by **L**_RW_) takes the form:

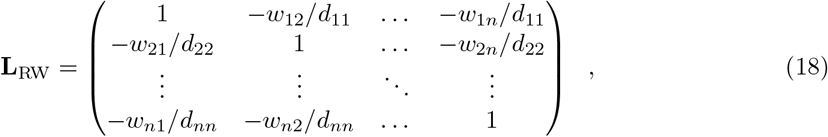

with 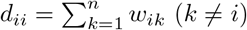 and *w*_*ij*_ = *w*_*ji*_. Eq. (16), then, becomes equivalent to 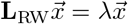, the standard eigenvalue equation for the Laplacian **L**_RW_ of a graph whose vertices all have degree 1. This new graph can be thought of as a derivative of the original, obtained by adjusting weights: if a vertex is connected to edges with large weights or if it has many neighbours – in other words, if it is strongly connected – the weights of the incident edges are scaled down. The opposite happens for a weakly connected vertex. Note that the random walk normalised Laplacian is not symmetric – indicating that the new graph is directed, i.e. the weights depend on the direction in which the edges are traversed. ^5^

The weight redistribution described above can be made more intuitive by analogy with people’s following on social media profiles, which may be regarded as a measure of their ‘degree of friendship’. Someone mainly interested in having a large following would likely accumulate a significant number of remote acquaintances among their connections. On the other hand, people who only connect with close friends would have a smaller following. In the former case, the ‘degree of friendship’ would have to be scaled down because a good proportion of connections would not represent meaningful friendships, while for the latter the percentage of followers who are intimate friends would be larger, and hence the degree of friendship should be scaled up.

In Ref. [34], the authors carry out tests on micro-array data and report that the generalized eigenvalue problem performs better at extracting information of biological interest. However, they focus on the extent to which the eigenvectors ^6^ (i.e. the vectors 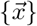 obtained by solving 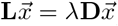) corresponding to the second and third smallest eigenvalues are able to reveal important features of the data by identifying relevant sub-clusters, whereas our interest lies in using the second smallest eigenvalue to infer how strongly connected a graph is. As will be shown in Section 3.2, we have found that by upping the effect of weakly connected vertices, the generalized eigenvalue problem tends to be more sensitive to noise. With this in mind, we shall henceforth focus on the standard eigenvalue problem. This is also the default method employed in the latest version of the VB toolbox.

### 2.2 The VB index as a modified cut-set weight

A connected graph (so called because any two vertices are joined by a path consisting of one or more edges) may be disconnected by removing edges. Let us suppose that given a connected graph *G*, we do away with a collection of edges (called a *cut* or *edge cut*) and manage to divide *G* into **two** components, *B* and *C*. The corresponding *cut-set weight* can be obtained by summing the weights of the ‘cut’ edges:

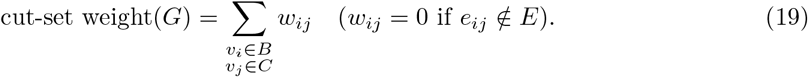

The rationale behind any clustering scheme is to group together vertices with strong affinity while separating those with divergent properties. In our case, the degree of similarity between any two vertices is reflected by the weight of the shared edge, and so we will attempt to partition the graph into two by removing the weakest edges. What we are interested in, therefore, is the cut that has the smallest weight.

To avoid instances when the cut-set weight is minimized by isolating a single vertex, we shall be using a modified cut-set weight (which we will henceforth refer to as the *ratio cut*) – one that takes into account the sizes of the resulting clusters: [32], [35], [37]:

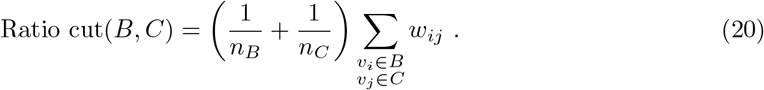

Here, *n*_*B*_ (*n*_*C*_) stands for the number of vertices in *B* (*C*).

The ratio cut can be expressed in terms of the graph Laplacian by means of Eq. (7). Let us consider a specific form for 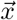 [35]:

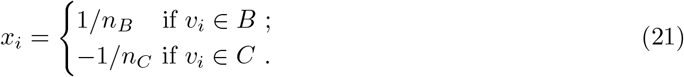

The magnitude (squared) of this vector is given by:

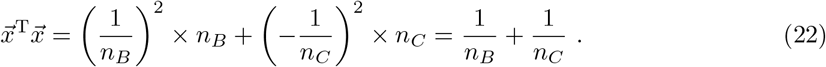

Substituting for *x*_*i*_ and *x*_*j*_ in Eq. 7 yields [35]:

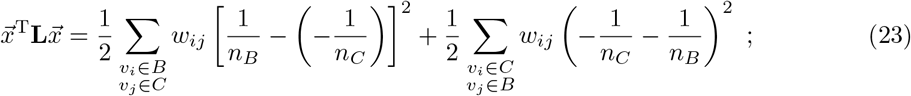

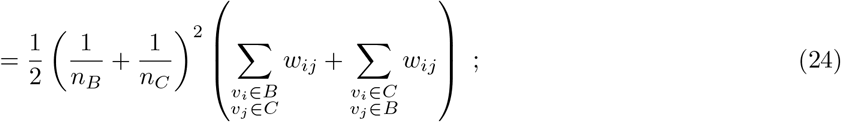

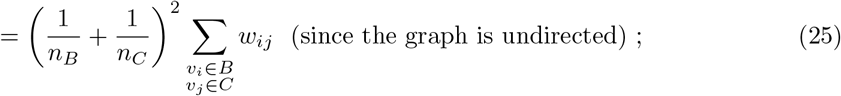

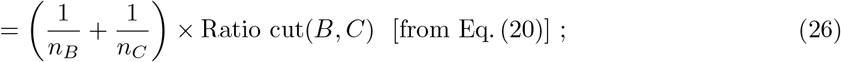

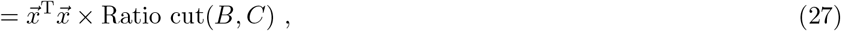

and hence

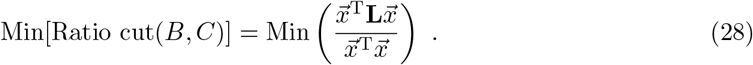

We note, however, that this holds provided 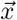 is as specified in Eq. (21), which in turn implies that 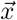 must be orthogonal to 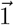, since [35]

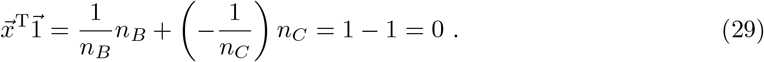

One common relaxation approach involves allowing the components of 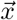 to take arbitrary values in the set of real numbers. Therefore, we now endeavour to find a vector 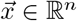 that minimizes the quantity 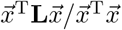 while still satisfying 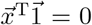. As shown in Section 2.1.2, this vector is none other than the Fiedler vector. We may consequently combine Eqs. (13) and (28) into one relation:

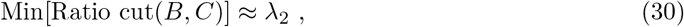

where the approximation sign reflects the fact that we have solved a relaxed version of Eq. (28).

In conclusion, we have shown that the second smallest eigenvalue of the graph Laplacian can be used to estimate a minimum value for the ratio cut, which is a sum of edge weights over all the edges removed to partition the graph into two clusters; this sum is weighted so that unbalanced clusters are penalized.

#### 2.2.1 Scaling the algebraic connectivity

Our next goal is to define a scaled version of λ_2_ that would be restricted to the range [0, 1]. It is well known that disconnected graphs have an algebraic connectivity of 0. At the other extreme, complete graphs (i.e. graphs in which every pair of vertices is connected via an edge) with maximally-weighted edges have the largest value of λ_2_, equal to the total number of vertices in the graph (so for a complete *n*-vertex graph whose edges all have a weight of 1, λ_2_ = *n*). We therefore scale the algebraic connectivity by the cardinality *n* of the vertex set (*n* is also called the *order* of the graph), and define the Vogt-Bailey (VB) index as follows:

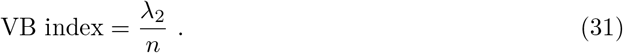

The difference between the scaling factor used here and the one in Ref. [1] stems from the fact that the original paper focuses on the generalized eigenvalue problem, 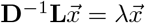, which in the case of a complete graph with maximally-weighted edges returns a value for λ_2_ equal to the mean of all eigenvalues except the smallest.

Substituting for λ_2_ using Eqs. (20) and (30) yields

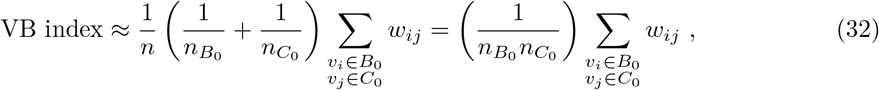

where a subscript 0 indicates that *B*_0_ and *C*_0_ are not just any two clusters, but the particular clusters that minimise the ratio cut, and the last equality was obtained by setting 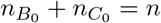. We shall henceforth refer to the cut-set weight 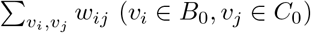 as the **VB cut**.

As expressed in Eq. (32), the VB index is extremely intuitive: it is the summed weight of the edges removed, divided by the total number of these edges. We emphasise that the VB cut corresponds to a special cut - the one that minimises the ratio cut. Secondly, 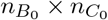 amounts to the number of edges dispensed with only if each vertex in cluster *B*_0_ is originally directly connected to every single vertex in *C*_0_. This essentially means that graphs which are not complete to start with are reinterpreted as complete graphs having some edge weights equal to zero. In other words, the meaning of the VB index is best understood if we consider the number of edges detached from the graph to be fixed at 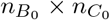, while the weights of those edges may vary – in the case of a complete graph with maximal weights, all edges have a weight of unity and as a result the VB index also equates to one, while a weakly-connected graph has edges with smaller weights (and possibly some with a weight of zero i.e. missing edges) and this lowers the VB index. It follows that a high VB index reflects the presence of ‘heavy’ edges and thus points to underlying voxels with strongly-correlated fMRI time series, while edges with small weights – due to weak correlations in said series – translate into a low VB value. Accordingly, a small VB index indicates a sharp change in local brain function.

## 3 Results

### 3.1 Testing the relation between the minimum ratio cut and the algebraic connectivity

To test how well the approximation of Eq. (30) holds, we assembled the following three sets of 27 *×* 27 affinity matrices:

- Set 1: 1061 matrices constructed from the functional MRI (fMRI) data for 2 participants (source: the Autism Brain Imaging Data Exchange (ABIDE) Preprocessed data set [38]).
- Set 2: 5074 matrices from the fMRI data for 10 participants (source: the minimally preprocessed Human Connectome Project (HCP) Young Adult data set [39]–[45]).
- Set 3: 118 matrices generated by sampling uniformly from the interval [0, 1].

Every matrix in Sets 1 and 2 corresponds to the neighbourhood graph of a randomly chosen vertex on the midthickness surface of the brain. For all 6253 matrices, we calculated λ_2_ as outlined in Ref. [1] (note, however, that the method of Ref. [1] is surface-based, whereas we employ the hybrid algorithm mentioned in the Introduction), and also determined the minimum ratio cut. The latter was computed by means of an exhaustive search over all possible 2-cluster partitions. The results are displayed in Fig. 1, from which it is immediately apparent that the relation given by Eq. (30) provides a good fit to the data, and compares well with the equation obtained via linear least-squares regression (*y* = 0.92*x−*0.08). The plot also reveals that matrices in Set 1 have, in general, a higher degree of connectivity than those in Set 2. This may be explained by the lower spatial resolution of the ABIDE data (3 mm isotropic, versus 2 mm isotropic for the HCP data). The manner in which the matrices in Set 3 were constructed makes it difficult to find a ‘fault line’ in the associated graphs, and consequently these graphs are harder to disconnect.

**Figure 1:**
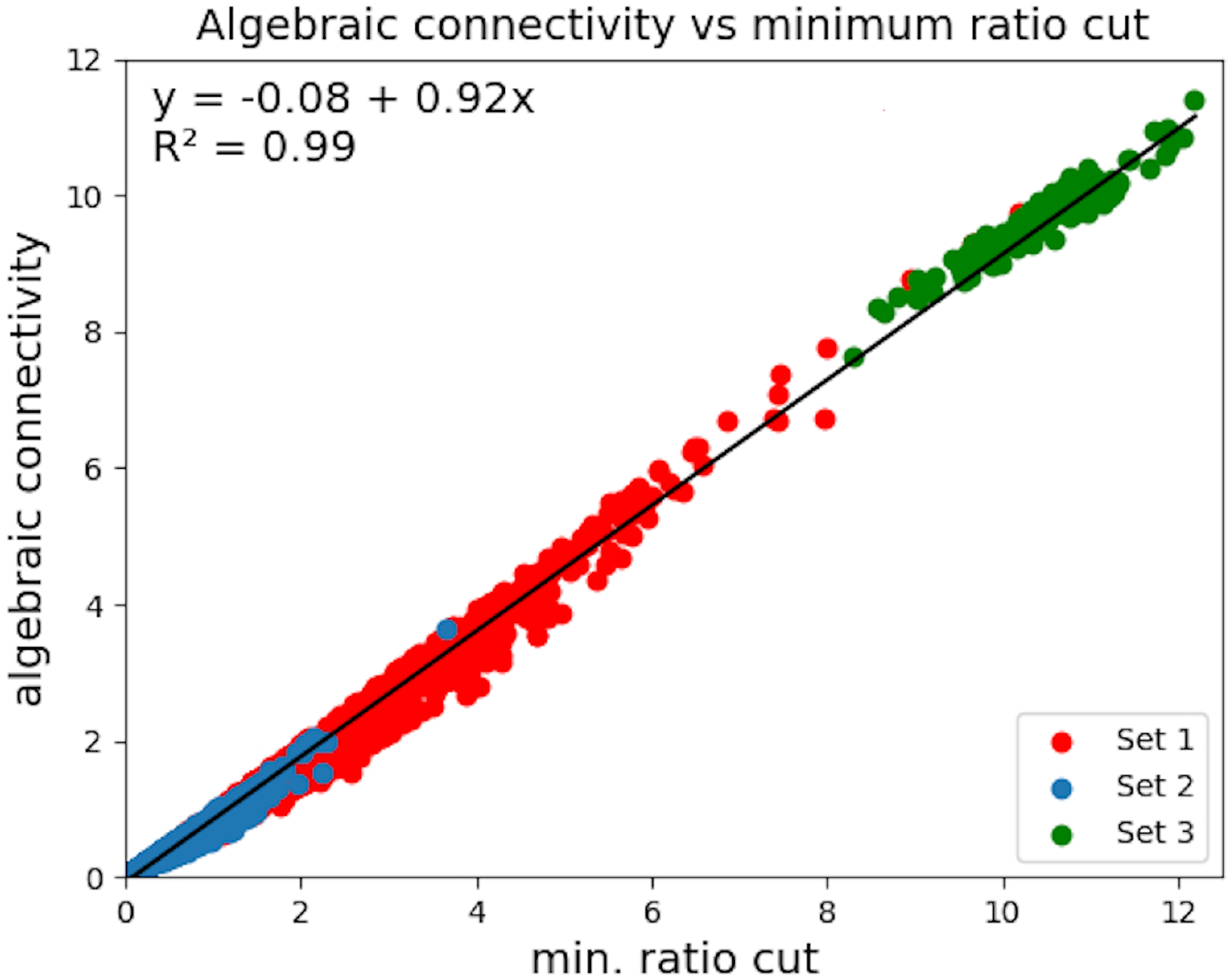
Algebraic connectivity vs minimum ratio cut. The former is equivalent to λ_2_, the second smallest eigenvalue of the graph Laplacian. The plot indicates that the minimum ratio cut is well approximated by the algebraic connectivity.

### 3.2 Comparison with ReHo

ReHo is arguably the method most commonly used in the literature to study local homogeneities in brain function [46]–[55]. In this section, we compare its performance with that of the VB index, the aim being to understand how these two measures differ in what they can tell us about activity in the brain.

The **Re**gional **Ho**mogeneity approach (ReHo) [46] employs Kendall’s coefficient of concordance (*W*) to gauge the degree of synchronisation between the time series of a general voxel and those of its nearest neighbours. *W* is given by [46]

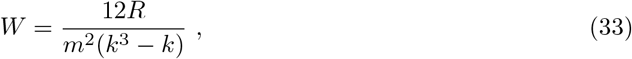

where m is the number of voxels in the neighbourhood, each with an associated time series of length *k*, and 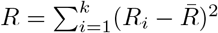. The quantity *R*_*i*_ is defined as the sum rank of the *i*^th^ data point. It is calculated as follows: we rank the *k* data points making up the time series of a given voxel, and repeat for all voxels in the neighbourhood. Then we sum the rankings of the *i*^th^ data point across the m voxels. 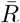 is simply the mean sum rank: 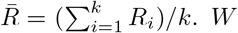 takes a value in the range [0, 1], with 1 indicating perfect synchronisation between the time series – i.e. a situation in which a given time point gets the same ranking for all voxels. A null value is obtained if the time series are completely out of sync.

ReHo is often used in combination with functional connectivity analysis to study whether certain disorders or diseases are associated with changes in local brain activity. Indeed, this approach has been adopted in the case of hepatitis B virus-related cirrhosis patients with or without minimal hepatic encephalopathy [47], as well as for patients with bipolar II disorder [48] or attenuated psychosis syndrome [49]. Additionally, abnormal ReHo values have been detected in subjects with early- or late-onset Parkinson’s disease [50] and acute or remitting multiple sclerosis [51], among others. ReHo maps can potentially serve as a non-invasive prognosis tool for cirrhotic patients with overt hepatic encephalopathy [52], and have a high diagnostic accuracy for congenital blindness [53]. Real-time fMRI neurofeedback and associated brain function self-regulation were found to impact the ReHo scores of brain regions involved in the processing of emotions [54]. In another study investigating the test-retest reliability of ReHo [55], the authors found that this could be improved by employing a fast imaging sequence, using nuisance correction but no spatial smoothing in the preprocessing stage, and by carrying out the analysis on the surface of the brain (in a vertex-wise manner, rather than the voxel-wise implementation of Ref. [46]).

The VB index differs from ReHo in three significant ways: firstly, it takes into account the values of the data points in the time series, not just their rankings. Secondly, a null score becomes highly improbable with ReHo if *k* is not equal to *m*, whereas the VB index is always zero for a disconnected graph. And thirdly, our framework admits a degree of flexibility – in that the similarity metric, which in our case is a modified Pearson correlation coefficient, can be replaced without compromising the inherent attributes of the VB index.

To compare the performance of the two local homogeneity measures, we consider 3 examples: a sparsely-connected graph [Fig. 2(a)], a complete graph [Fig. 2(b)] and a disconnected one [Fig. 2(c)]. These graphs were constructed by assigning a time series ^7^ of length 20 to each vertex and calculating the affinity matrix as detailed in Ref. [1]. The entry *a*_*ij*_ in the affinity matrix, derived from the correlation between the time series of vertices *υ*_*i*_ and *υ*_*j*_, then serves as the weight of the edge joining the two vertices. We generated the data in such a way that if the time series of a vertex is thought of as a vector, two vertices not directly linked by an edge would have orthogonal time series, which in turn would imply an edge with essentially zero weight. In the three cases represented in Fig. 2, the ReHo value turns out to be higher than the VB index. **It is especially interesting to note that while the latter is approximately zero for the disconnected graph, and varies by an order of magnitude between the sparsely connected and complete graphs, the values we get with ReHo are comparable across all three examples**.

**Figure 2:**
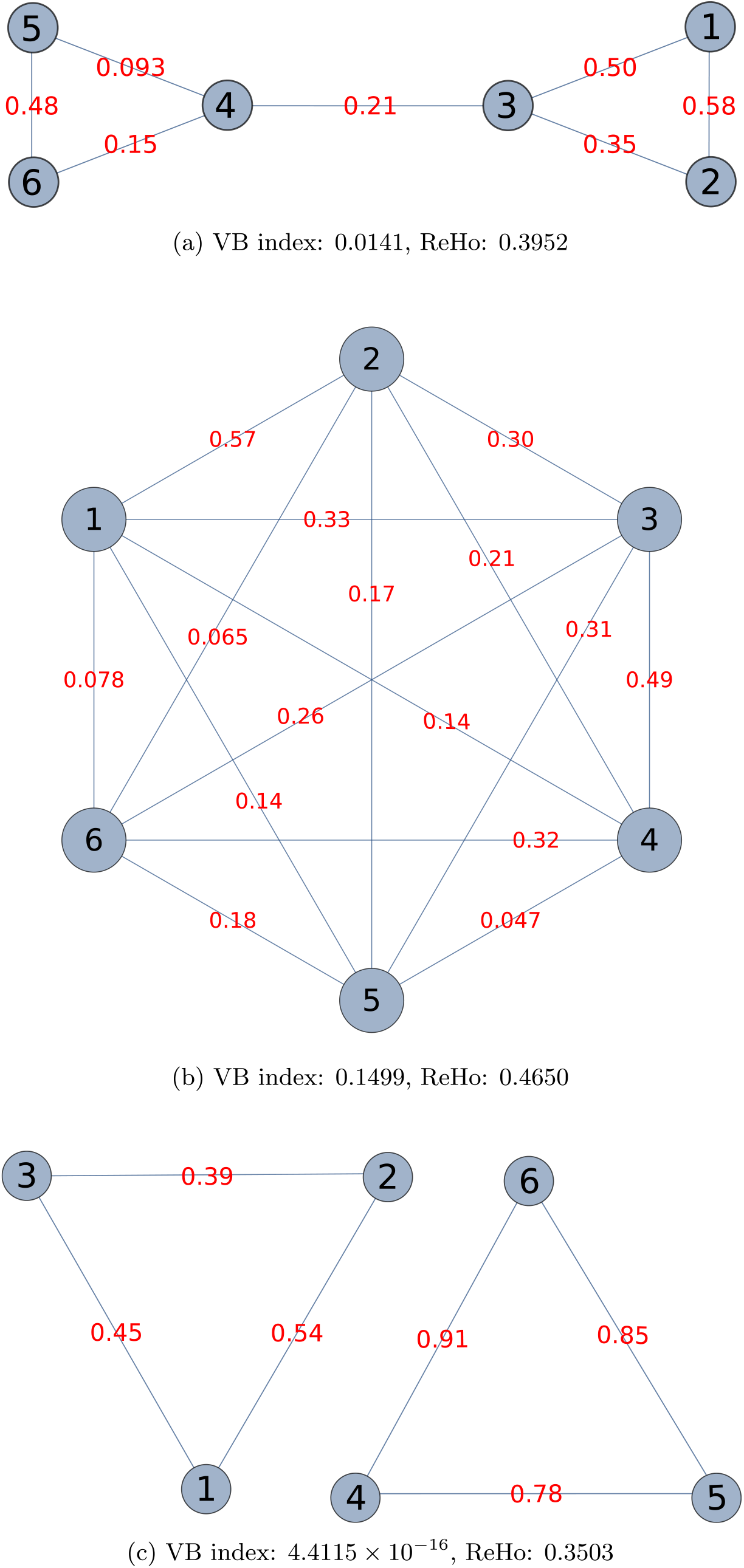
Different ways of joining 6 vertices into a graph. The examples shown are (a) sparsely connected, (b) fully connected (complete) and (c) disconnected.

Next, we tested ReHo and the VB index using synthetic fMRI data produced with the R software package `neuRosim` [56]. The data had a signal-to-noise ratio of 3 and consisted of 5 spherical task activations with hard edges, superimposed on a mixture of the following noise components [56]:

- **white** – a Rician distribution with non-centrality parameter of 0 (5%);
- **temporal** – an autoregressive model of order 3 (10%);
- **low frequency drift** – the frequency was set to 128 s (1%);
- **physiological** – noise due to heart beat and respiration (9%);
- **task** – noise due to spontaneous neural activity at the activation sites (5%);
- **spatial** – a Gaussian random field generated by a kernel having a full width at half maximum of 4 (70%).

The synthetic data were processed by a hybrid algorithm which maps a given vertex on the midthickness surface of the brain to the corresponding voxel. It then calculates the ReHo or VB index value for a 27-voxel neighbourhood centred around (and including) the principal voxel, and maps the result back to the original vertex. The algorithm proceeds this way in a vertex-wise manner and at the end returns a VB or ReHo map for the entire brain surface. The resting-state data used to generate a baseline image and volumetric mask for the construction of synthetic time series (the volumetric mask is also required to run the version of the toolbox employed for this paper), as well as the midthickness surface and cortical mask provided as input to the hybrid algorithm and the inflated surface needed to generate the figures, were obtained from the minimally preprocessed HCP Young Adult data set [39]–[45] by randomly selecting one participant.

The outputted VB and ReHo brain maps are presented in Figs. 3 and 4, respectively. It can be seen that both algorithms easily identify the activations, and as anticipated, produce an adequate level of contrast between them and the underlying noise. However, the activated regions have sharper edges in the VB map, and the accompanying histograms reflect this clearly. In the case of the VB index, that part of the histogram at the upper half of the range of VB values forms separate clusters that directly correspond to the areas of activation, but the histogram for the ReHo metric does not share this feature.

**Figure 3:**
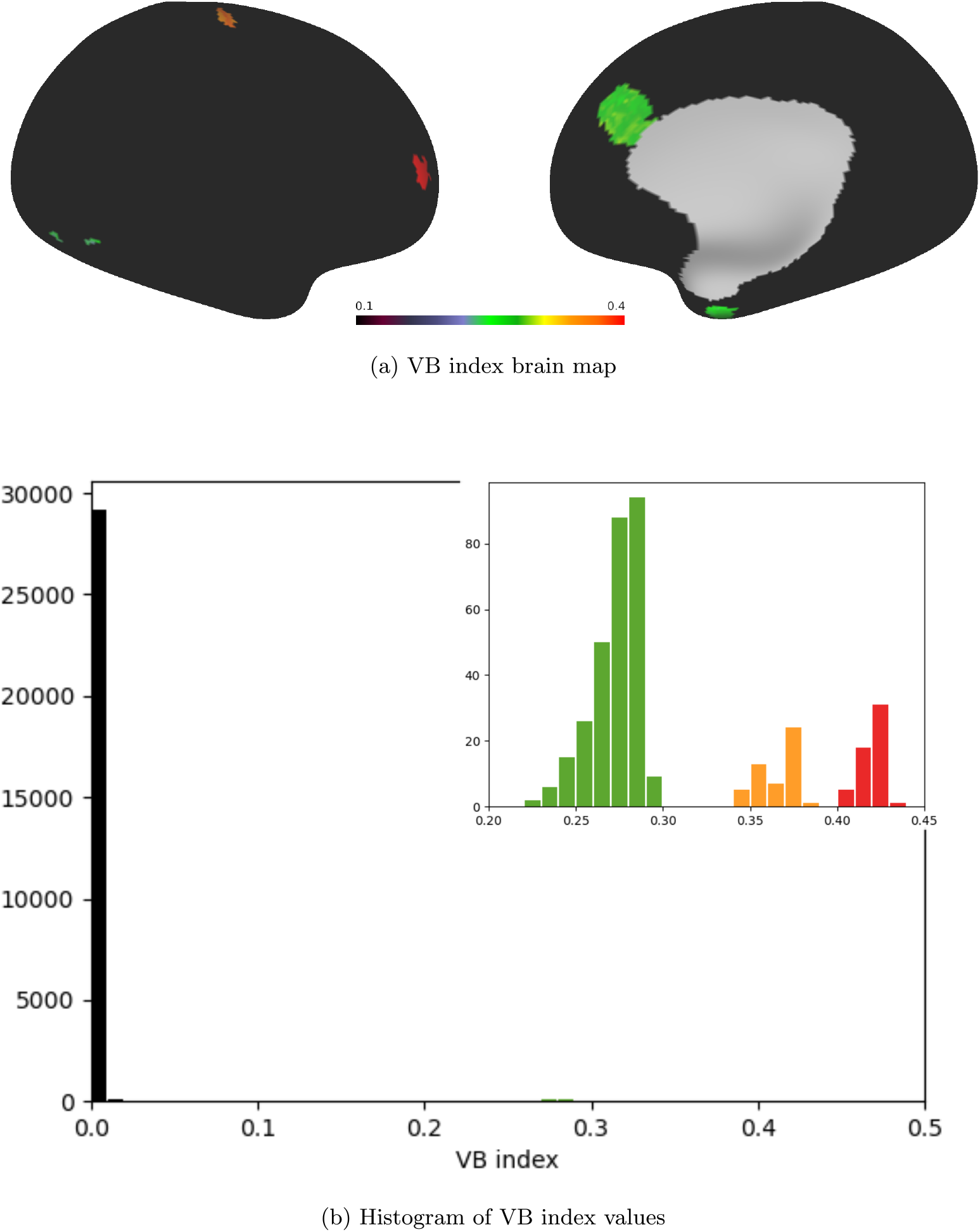
Brain map of VB index values (top) and associated histogram (bottom). The greater majority of the synthetic data provided to the algorithm represent noise and have a VB index close to zero. The inset shows 3 clusters at higher VB values. These clusters arise due to the task activations superimposed on the noise and have colour correspondence with the activated regions in the top panel.

**Figure 4:**
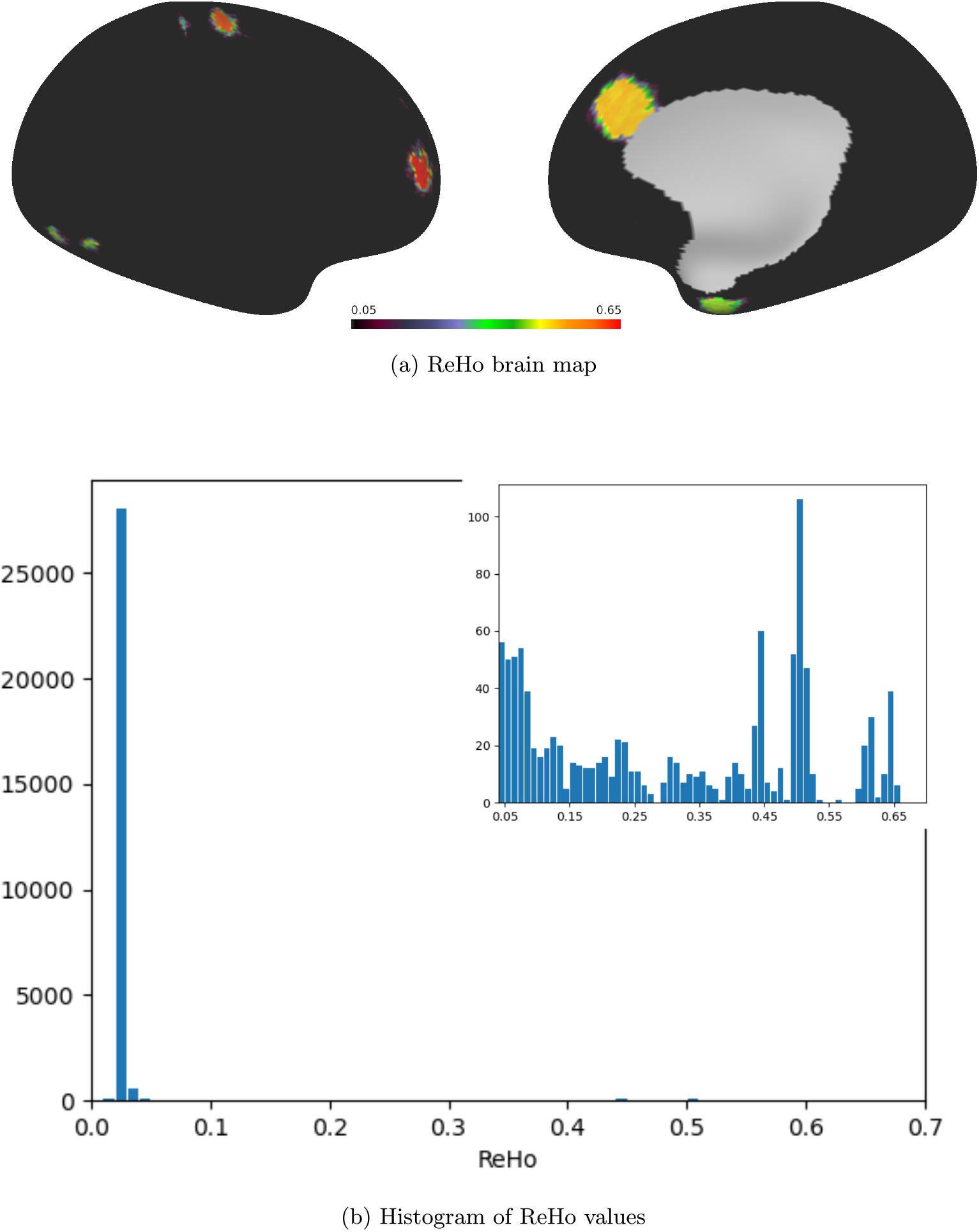
Brain map of ReHo values (top) and associated histogram (bottom; the inset displays data binned over a sub-interval). Like the VB index, the ReHo metric clearly differentiates between the activated regions and the background noise, but the distinction among the activations themselves is significantly sharper in the case of the VB index.

The same synthetic data were also fed into the VB toolbox with the Laplacian normalization set to geig, which instructs the algorithm to return the eigenvalues of the generalized eigenvalue problem [Eq. (16)]. Fig. 5 shows the distribution of VB index values obtained, both as a brain map and as a histogram. It is immediately apparent that the `geig` method distinguishes much less sharply among the individual activated regions than the original approach (`unnorm`), which, we recall, is based on the standard eigenvalue problem 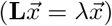. Additionally, the VB indices that geig outputs for the noise component have higher values and greater variance than their `unnorm` counterparts, indicating more sensitivity to noise.

**Figure 5:**
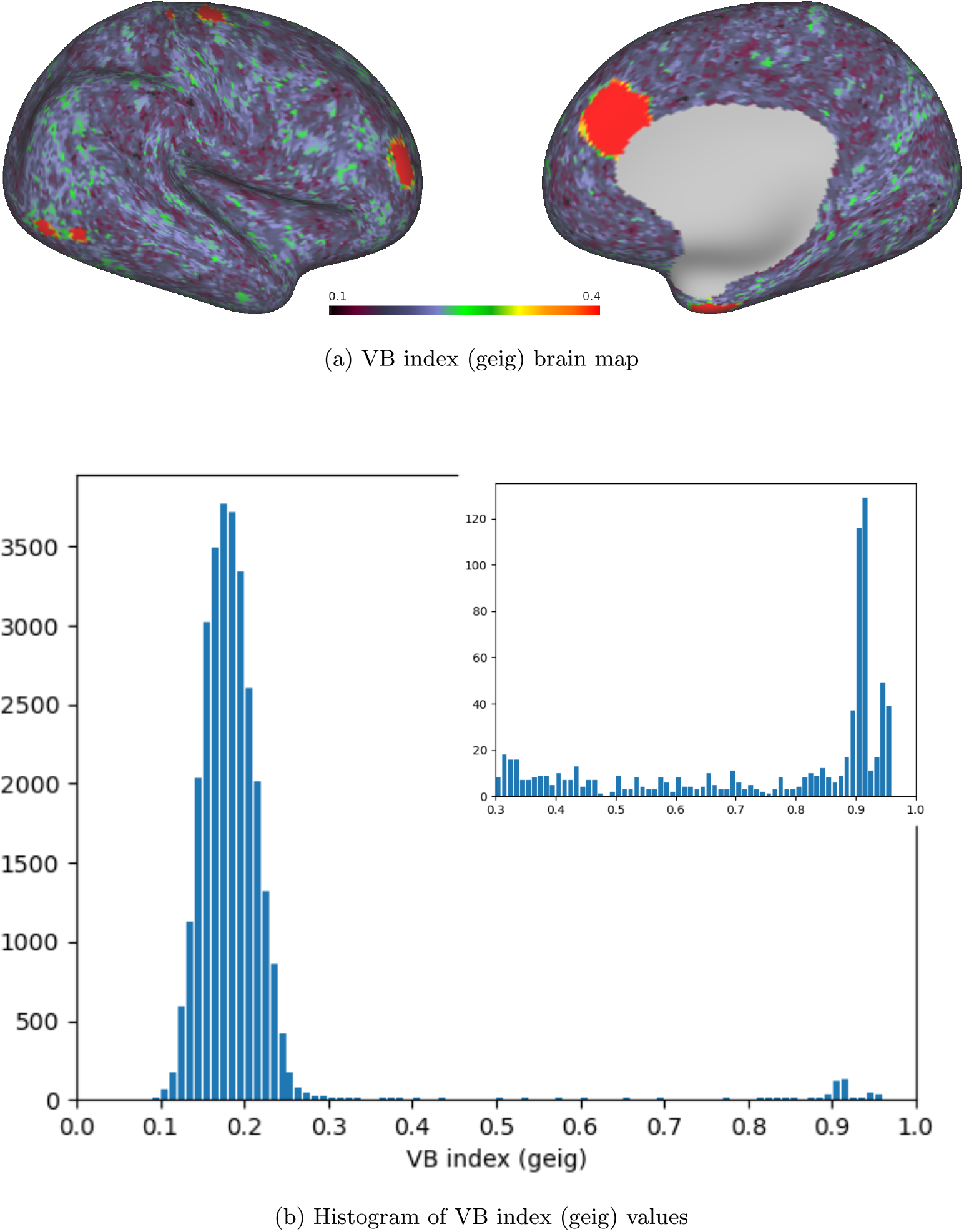
Brain map and histogram for the VB values obtained with the `geig` method. The inset presents the data binned over a sub-interval to the right of the principal peak.

## 4 Discussion

The VB index is an ‘edge-detection algorithm’ introduced in Ref. [1] to look for sharp changes (‘edges’) in the local functional organization of the human cortex. In this work, we expound on the details of the underlying mathematical framework. In particular, we re-interpret the VB index as a modified cut-set weight associated with a particular graph cut – one that finds a trade-off between eliminating as few edges as possible and having clusters with a comparable number of vertices. This makes the VB index extremely intuitive. We test the approximation on which our interpretation is based using matrices extracted from real data, and conclude that it holds very well. Additionally, we introduce the concept of a VB cut (which is simply the sum of edge weights associated with the cut mentioned above), and show that the VB index can be understood as the VB cut divided by the total number of edges removed to partition the graph into two (provided missing edges are treated as edges with null weight).

Next, we compare the performance of the VB index with that of ReHo, a metric commonly used to assess regional homogeneity in brain function. The two algorithms can be executed with a similar amount of computational effort. We apply them to synthetic functional MRI data generated with the `neuRosim` package [56], and plot histograms for the output. While both ReHo and the VB index pick out the areas of activation from the background noise, the VB index traces sharper borders around these areas, localizing the activations with greater precision. We also investigate whether solving the generalized eigenvalue problem – rather than the standard one – to calculate the VB index has any benefits. Our results show that when determined this way, the VB index is more sensitive to noise and does not distinguish as well among the different regions of activation. Consequently, new versions of the VB toolbox will, by default, employ the regular Laplacian and the standard eigenvalue problem.

## Data availability

Data were provided in part by the Human Connectome Project, WU-Minn Consortium (Principal Investigators: David Van Essen and Kamil Ugurbil; 1U54MH091657) funded by the 16 NIH Institutes and Centers that support the NIH Blueprint for Neuroscience Research; and by the McDonnell Center for Systems Neuroscience at Washington University.

All research data will be made available upon publication.

## Code availability

The version of the VB toolbox employed in this work may be obtained from GitHub (https://github.com/VBIndex/py_vb_toolbox/tree/Local-gradients-paper). Installation instructions are provided in the README file. Kindly note that the Laplacian normalization method still defaults to `geig` in this version.

The brain maps in Figs. 3–5 were prepared with the Connectome Workbench software, v1.4.2 [57], while the graphs in Fig. 2 were generated with Wolfram Mathematica v13.0.1.0 [58].

## Acknowledgements

This article was produced as part of the BEyond BOundaries on the Brain (BE-BOB) Project, financed by the University of Malta Internal Research Grants Programme (Research Excellence Fund – grant agreement no. 202201). It is based upon work from COST Action CA18106 - The neural architecture of consciousness (NeuralArchCon), supported by COST (European Cooperation in Science and Technology: www.cost.eu). Preliminary results were outlined in the poster entitled *VB Cut: a measure of local correlations in brain function*, which was presented at OHBM 2022. The authors would like to thank Dr L. Q. Costa Campos for contributions to the software.

The research reported on in this article was conducted in accordance with the review procedures of the University of Malta’s Research Ethics Committee (UREC).

## CRediT statement

**CF:** Conceptualization, Methodology, Formal analysis, Writing - original draft preparation, Writing - review & editing; **PG:** Software, Writing - review & editing; **IAI:** Writing - original draft preparation, Writing - review & editing; **KS:** Conceptualization, Methodology, Writing - review & editing; **CJB:** Conceptualization, Methodology, Writing - review & editing.

## Footnotes

1 The larger the number of edges and the weights assigned to them, the more strongly-connected the graph is, and the greater the challenge to break it up into separate parts.

2 The proof is provided with the supplementary data of the companion paper, Ref. [1]. A superscript T indicates that a quantity is transposed.

3 This constraint may be relaxed in the manner of Ref. [34].

4 Since **L** is a symmetric matrix, it has a full set of linearly independent (mutually orthogonal) eigenvectors.

5 The equation 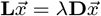 may also be cast in the form 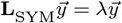, where **L**_SYM_, the *symmetric normalised Laplacian*, is defined as **D**^*−*1*/*2^**LD**^*−*1*/*2^, and 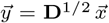 [1] (equivalently, 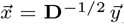). In this case, the edges would remain undirected. Note that **D**^*−*1*/*2^ can be determined from **D** by finding the reciprocal of the elements along the main diagonal of **D** and computing their square root.

6 Strictly speaking, the authors work with the symmetric normalised Laplacian and adopt 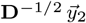 as the normalised Fiedler vector (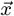 and 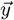 have the same meaning as in Footnote 5, and 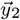 is the eigenvector corresponding to the second smallest eigenvalue of **L**_SYM_).

7 The time series were each modelled as a sine wave with a noise component drawn randomly from a normal distribution. Distinct series differed in the initial phase of the sine function, as well as in the mean and variance of the normal distribution.

